# GWAC: A machine learning method to identify functional variants in data-constrained species

**DOI:** 10.1101/2024.11.15.623873

**Authors:** Andrew G. Sharo

## Abstract

As environments change, the ability of species to adapt depends on the functional variation they harbor. Identifying these functional variants is an important challenge in conservation genetics. Due to the limited data available for most species of conservation interest, genome-wide selection scans that link specific genetic variants with a phenotype are not feasible. However, functional variants may still be identified by considering predicted consequence, evolutionary conservation, and other sequence-based features. We developed Genome-Wide vAriant Classification (GWAC), a supervised machine learning framework to prioritize genome-wide variants by functional impact. GWAC requires only features that can be generated from an annotated genome. We evaluate GWAC by first using a set of human data constrained to match what may be available for threatened species. We find that GWAC weights features more heavily that are known to be predictive of functional variation and prioritizes both single nucleotide variants and indels, consistent with mutational constraint found in population genetics studies. GWAC performs nearly as well as CADD, a leading genome-wide predictor in humans that uses substantially more features and data that are typically available only for model organisms. While it is not possible to empirically evaluate GWAC on a species for which no functional variants are known, we find that a version of GWAC generated for the greater prairie chicken (*Tympanuchus cupido pinnatus*) weights features similarly to our human version. We compare the results of using a species-specific variant impact predictor against lifting-over variants from a closely related model organism and find that the species-specific approach retains functional variants that are lost during lift-over. We anticipate GWAC could be used to estimate conservation metrics such as genetic load and adaptive capacity, while also enabling researchers to identify individual variants responsible for adaptive phenotypes.

## Introduction

Researchers can better assess and conserve threatened species when aided by genomics (1-3). Whole genome sequencing enables researchers to measure key indicators of species health including genetic diversity, adaptive potential, and genetic load (4). These indicators benefit when functional variants can be distinguished from broader neutral genetic diversity. Functional variants impact phenotype appreciably and can be adaptive or deleterious, in contrast with neutral variants. Indeed, there is growing appreciation that functional diversity predicts species health better than bulk genetic diversity (5). Additionally, in applications of assisted evolution or de-extinction, it is valuable to know which functional genetic differences between species may result in their distinct phenotypes.

Functional variants are challenging to identify in species with limited genetic diversity and that are rare or difficult to sample or phenotype (3). These limitations impede traditional approaches such as genome-wide selection scans that link specific genetic variants with a phenotype. Such scans require data sets of sizes that are often unobtainable from threatened species. As an alternative approach, PhyloG2P methods associate shared phenotypes across a phylogeny with genomic regions (6). However, PhyloG2P requires that several species have the same phenotype of interest. Additionally, this approach typically identifies genetic regions associated with phenotypes across species, rather than pinpointing specific functional variants.

Instead of associating variants with a phenotype, an alternative technique to identify functional variants is to consider the genomic context of variants. This approach has been enabled by the recent growth of reference genomes for threatened species (7, 8). With an annotated reference genome, functional diversity can be assessed by considering nonsynonymous variation (e.g., McDonald-Kreitman test (9)), and individual variants can be predicted to be functional using SIFT scores (10). However, these methods do not address functional genetic diversity outside of nonsynonymous variants. Noncoding regions can also now be evaluated thanks to the growing availability of reference genomes and multiple genome alignments. With multiple genome alignments, researchers can identify conserved or rapidly evolving regions of genomes using approaches such as phastCons (11), phyloP (12), and GERP (13). Additionally, tools such as SnpEff (14) and VEP (15) can predict the consequence of variants (e.g., frameshift, splice acceptor) based on a genome annotation.

While some of these approaches prioritize variants genome-wide by function, none alone is complete. Variants with a “High” SnpEff impact often serve as an indicator for deleteriousness (16-19). However, this is a blunt category that almost certainly includes neutral variants. For example, SnpEff categorizes all stop-gain variants as “High” impact, but we know that the average human genome contains more than a dozen rare benign stop-gain variants (20). phastCons and phyloP can similarly score variants genome-wide, however researchers have found that a combination of sequence features with conservation scores consistently outperforms conservation scores alone in predicting functional variants in mouse and human (21).

Prediction of functional variants in genomes is improved by integrating multiple genomic features simultaneously using a machine learning approach. Such methods are generally referred to as variant impact predictors (22). One popular variant impact predictor in humans is CADD (23), which predicts the functional importance of variants genome-wide. CADD considers more than 100 genome-wide features and combines them into a single score. In addition to evolutionary conservation/acceleration and variant consequence, CADD features also include experimentally derived features such as chromatin accessibility, histone modifications, and transcription factor binding sites. Similar predictors following the CADD framework have been created for mouse (21), pig (24), and chicken (25). However, these genome-wide variant impact predictors are thus far limited to only data-rich model organisms. For some non-model organisms, it may be possible to lift-over CADD scores from closely related model organisms (e.g., from humans to chimps), however this approach may result in lower accuracy (21).

Here, we investigate whether the limited genomic features available for most threatened species are sufficient to create species-specific predictors to identify functional variants genome-wide. We developed a supervised logistic regression predictor, GWAC (Genome-Wide vAriant Classification). We trained GWAC to distinguish a dataset enriched for neutral variants from a dataset enriched for functional variants. GWAC combines the frameworks of GWAVA (22) and CADD. Like GWAVA, GWAC uses common variants in a population to generate neutral-enriched training data. Like CADD, GWAC simulates variants to create functional-enriched training data. The GWAC framework has important advantages over CADD when data-limited species are considered. CADD trains on a neutral-enriched dataset consisting of variants that are derived compared to an inferred ancestral genome (last common ancestor with chimpanzee) and fixed or nearly fixed (AF > 95%) in humans. The CADD framework has the following drawbacks that make it difficult to implement for threatened species: 1) CADD requires allele frequencies to be known with enough precision to determine if derived variants are fixed or nearly fixed, which entails sequencing many individuals from a population. 2) A predictor trained using the CADD framework will be less capable of identifying functional variants between closely related species, as many of these variants are likely to be derived and thus included in the neutral-enriched training set, biasing predictions at these positions. 3) Finally, the CADD framework requires identifying an appropriate related species with which to infer an ancestral genome, which may be challenging for species in poorly sequenced taxa. In contrast, GWAC resolves these drawbacks, as common variants can more easily be identified from a small number of genomes, they do not include fixed differences between species, and an inferred recent ancestral genome is not required.

We find that the limited features typically available for threatened species are capable of accurately identifying functional variants genome-wide. The GWAC framework learns features of functional variants that are consistent with our biological understanding of functional variation. A version of GWAC trained on constrained human data distinguishes known pathogenic variants from common variants nearly as well as the data-rich predictor CADD. A version of GWAC created for a data-limited species, the greater prairie chicken, appears comparable with our human predictor. Finally, we compare our species-specific approach to an approach that lifts-over predictions from a related model organism.

## Results

### We designed GWAC for data-limited species

We implemented GWAC as a logistic regression classifier with 47 features (see Methods). These features included evolutionary conservation/acceleration (phastCons, phyloP, and SIFT), variant consequence (e.g., nonsynonymous), genomic context (GC content, CpG content), and variant attributes (e.g., alternative allele, variant length). We chose these features because they can be generated for most species with a reference genome. To score a variant of interest, GWAC calculates a weighted linear combination of the features associated with the variant, which is then fed into a logistic model. These weights reflect the predictive value of each feature and are learned during training.

We trained GWAC to distinguish a dataset enriched for neutral variants from a dataset enriched for functional variants. We created the neutral-enriched dataset by gathering all variants found in three randomly selected individuals from the 1000 Genomes Project (1KGP), a diverse collection of human genomes. Since most of these variants have been acted upon by selection, they should be enriched for neutral variants. We created the functional-enriched dataset by randomly generating variants (see Methods). Since these random variants have not been acted upon by selection, they should be more enriched for functional variants than the neutral-enriched dataset. In total, more than 15 million variants were used to train our human version of GWAC.

### GWAC learns general properties of functional variants

GWAC learned that evolutionary conservation/acceleration scores are particularly predictive of functional variants (Fig. 1A). The logistic regression model underlying GWAC assigns larger weights to features that it learns are more predictive. GWAC learned that the most predictive feature by far was the evolutionary conservation/acceleration score PhyloP of a 7 vertebrate alignment. This alignment consists of the genomes of three primates, two rodents, dog, and opossum. This large weight suggests that phyloP alignments of a relatively small group of related species are particularly predictive of functional variants, consistent with existing predictors in model organisms (21, 26). Secondary predictive features included additional evolutionary conservation/acceleration scores, certain variant consequences (e.g., nonsynonymous), variant attributes (e.g., indel length), and GC content. Gene expression, CpG sites, and several variant consequences were learned by GWAC not to be particularly important.

**Fig. 1.**
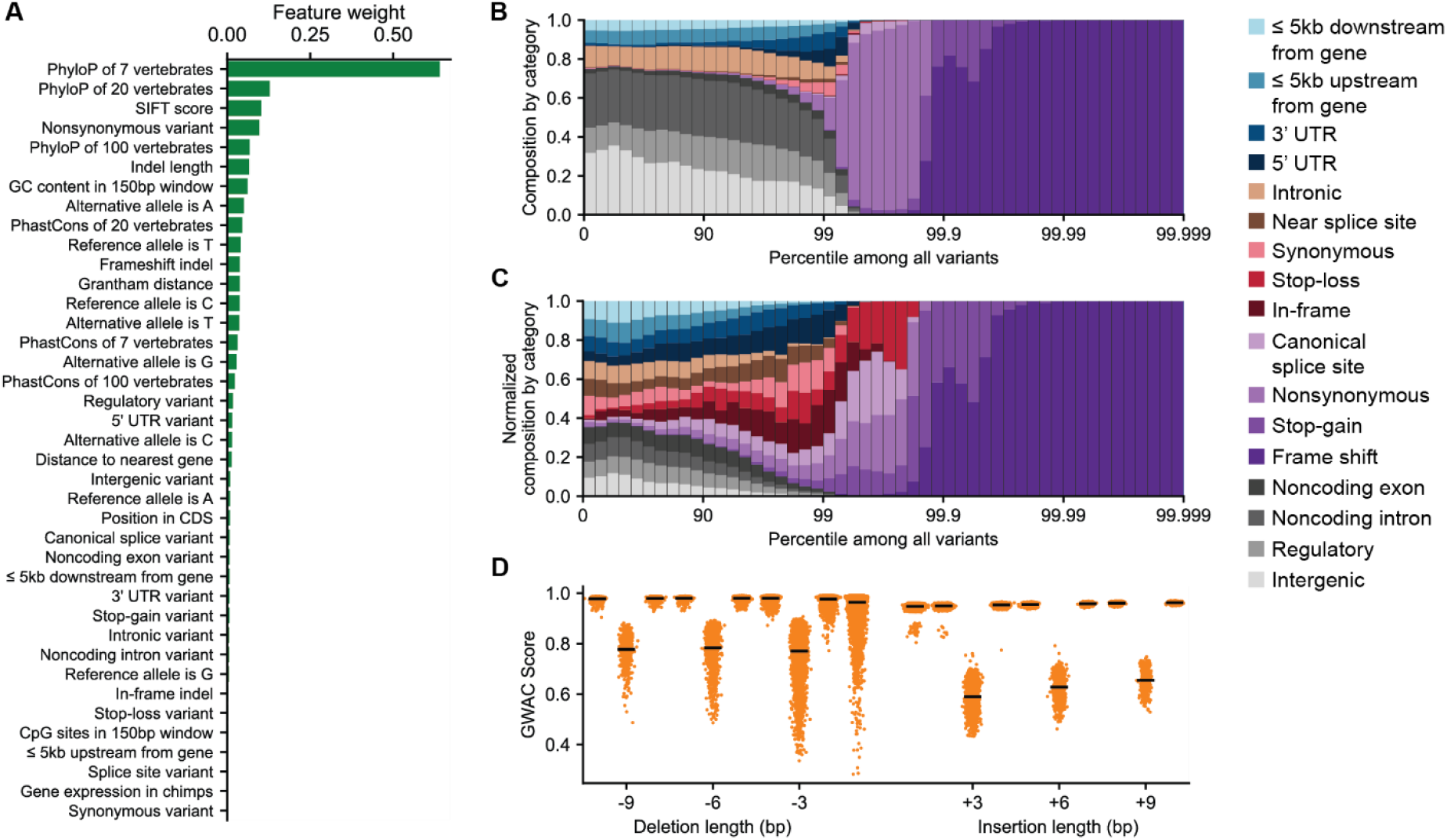
GWAC identifies general properties of functional variants in humans. **A**) Absolute values of feature weights learned by GWAC. **B**) Composition of variant categories in each GWAC score percentile bin. 10 million variants were randomly generated to create this figure. Detailed descriptions of each consequence are provided in Table 1. **C**) Equivalent data as B, with variant categories normalized by the number of variants in each category. **D**) GWAC scores of frameshift and in-frame indels randomly generated in the human genome. Black bars indicate mean values. Larger GWAC scores indicate greater chance of being functional.

GWAC ranks variants consistent with our knowledge that certain variants consequences are more likely to lead to a functional effect than others (27). We generated 10 million random variants and asked GWAC to score these variants. We found that variants in the 99th percentile of scores or greater (top 100,000 functional variants of 10 million total variants) were comprised largely of frameshift, stop-gain, non-synonymous, and canonical splice site variants (Fig. 1B,C). Additionally, there are also a small number of synonymous, intronic, and even intergenic variants in the 99^th^ percentile or above, which is in agreement with studies that have found variants in these categories can have functional impact, although rarely (28-30).

**Table 1.** Detailed description of all features used for training predictors. Note that Distance, phastCons7, phastCons20, phastCons100, phyloP7, phyloP20, and phyloP100, CDS_position also had Boolean indicator variables when they were absent.

| Name | Type | Description |
| --- | --- | --- |
| Ref | String | One hot encoding of reference allele |
| Alt | String | One hot encoding of alternative allele |
| Indel Length | Int | Negative for deletions, positive for insertions, zero for SNVs |
| Consequence | String | One hot encoding of simplified VEP consequence (5PRIME_UTR, CANONICAL_SPLICE, DOWNSTREAM, FARME_SHIFT, INFRAME, INTERGENIC, INTRONIC, NONCODING_CHANGE, NONCODING_EXON_CHANGE, NON_SYNONYMOUS, REGULATORY, SPLICE_SITE, STOP_GAINED, STOP_LOST, SYNONYMOUS, UPSTREAM) |
| Distance | Int | Distance to closest gene (5kbp max) |
| CDS Position | Int | Position within coding sequence, where first nucleotide is 0 |
| phastCons7 | Float | phastCons score from 7 vertebrate alignment |
| phastCons20 | Float | phastCons score from 20 vertebrate alignment |
| phastCons100 | Float | phastCons score from 100 vertebrate alignment |
| phyloP7 | Float | phyloP score from 7 vertebrate alignment |
| phyloP20 | Float | phyloP score from 20 vertebrate alignment |
| phyloP100 | Float | phyloP score from 100 vertebrate alignment |
| GC | Float | GC content of surrounding +/- 75bp |
| CpG | Float | CpG content of surrounding +/- 75bp |
| Grantham | Int | Grantham distance between ref and alt amino acid |
| Expression | Float | Maximum gene expression of all overlapping genes |
| SIFT | Float | SIFT score |

GWAC learned properties of functional indels that are consistent with findings from human population genetics. From population studies, researchers have found that frameshift variants face greater negative selection than in-frame indels (22). GWAC ranks frameshift variants as significantly more functionally important than in-frame indels when asked to rank random variants (Fig. 1D). Additionally, GWAC ranks deletions, particularly in-frame deletions, as more functionally important than insertions. This distinction is consistent with research in humans finding deletions are generally more deleterious than insertions (31).

### With limited data, GWAC nearly matches the performance of CADD, a data-rich predictor

To assess GWAC’s ability to distinguish functional from neutral variants, we compared it with several existing predictors. The first is CADD, one of the most-cited variant impact predictors in human clinical genetics (22). In addition to all the features that GWAC considers, CADD also considers dozens of extra features, including those that capture chromatin accessibility, histone modifications, and transcription factor binding sites derived from experimental data. These features are typically unavailable for species of conservation interest. Additionally, CADD uses a distinct training data framework (described above). We also compare GWAC to a CADD-limited predictor that uses the same training framework as CADD, but only includes the features used by GWAC. This allows us to directly compare the GWAC and CADD training frameworks. Finally, we compare with SnpEff impact (‘High’, ‘Moderate’, ‘Low’, or ‘Modifier’), which is commonly used to prioritize variants found in data-limited species (16-19).

We assessed performance by evaluating how well these methods distinguished pathogenic from common variants in humans. Pathogenic variants were collected from ClinVar, a catalog of clinically relevant variants, and common variants were collected from 1KGP. Pathogenic and common variants at the same position as variants used to train GWAC or CADD were removed from consideration. To avoid training and testing on nearby variants, GWAC and CADD-limited were evaluated using a leave-one-chromosome out approach (see Methods). We split test variants into 3 non-overlapping groups: coding (any alteration to CDS), splicing (canonical and splice-region variants), and non-coding (all other variants). We measured performance using the area under the receiver-operating characteristic curve (AUC).

When distinguishing pathogenic from common variants, GWAC performed substantially better than or equivalent to CADD-limited and SnpEff, and nearly as well as CADD. Among coding variants (Fig. 2A), GWAC (AUC=0.96) and CADD (AUC=0.96) performed similarly well, with SnpEff (AUC=0.92) and CADD-limited (AUC=0.90) performing worse but still strongly. Among splicing variants (Fig. 2B), CADD (AUC=0.99) performed exceptionally, with GWAC (AUC=0.97), SnpEff (AUC=0.97), and CADD-limited (AUC=0.96) performing similarly well. Non-coding variants were significantly more challenging to classify (Fig. 2C), and there was no significant difference between GWAC, CADD, and CADD-limited (all AUC=0.76), with SnpEff (AUC=0.51) performing no better than random. SnpEff performed poorly because it annotated nearly all non-coding variants with ‘Modifier,’ its lowest impact annotation. Finally, we evaluated these predictors on only missense variants (Fig. 2D), and compared them to two additional predictors, SIFT and PROVEAN, that are designed to predict the impact of missense variants and are commonly used in non-model organisms (32-36). CADD (AUC=0.92) performed best, followed by CADD-limited (AUC=0.90), GWAC (AUC=0.88), PROVEAN (AUC=0.87), and SIFT (AUC=0.86). Finally, SnpEff performed only slightly better than random (AUC=0.53), as it annotated the vast majority of missense variants as ‘Moderate’. Since GWAC is trained to distinguish functional from common variants, we expect it to do well in this scenario. Yet, when these analyses were repeated on ability to distinguish rare pathogenic from rare benign variants, we observed a similar trend (Fig. S1).

**Fig. 2.**
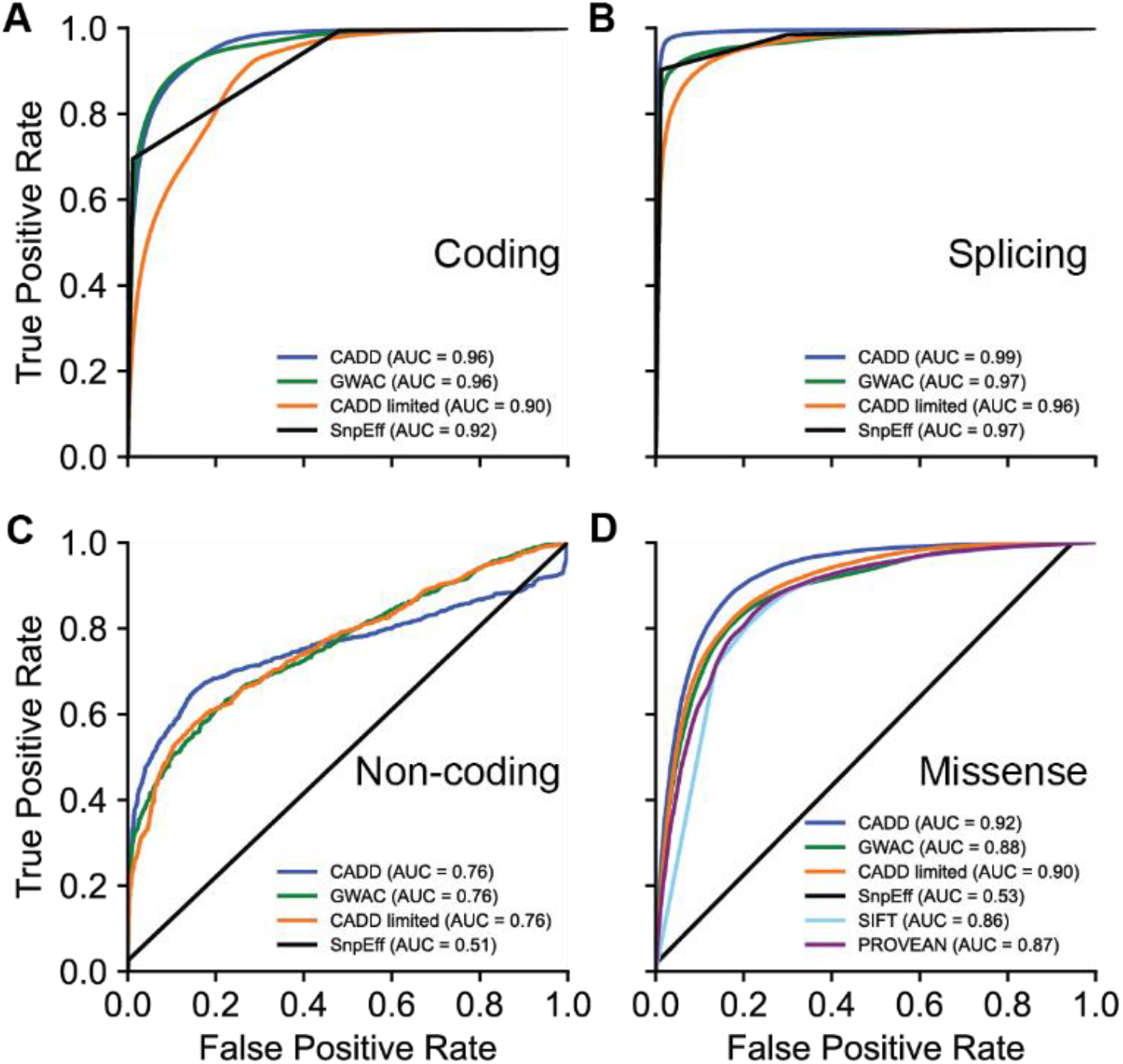
GWAC nearly matches the performance of data-rich predictor CADD, while outperforming existing predictors for non-model organisms. Receiver-operating characteristic (ROC) comparison of GWAC with other genome-wide predictors on ability to distinguish pathogenic from common variants in **A**) coding regions **B**) splicing regions, and **C**) non-coding regions. **D**) ROC comparison of GWAC with missense predictors.

### Our greater prairie chicken GWAC learns properties of functional variants consistent with the human GWAC

We next created a version of GWAC for a data-limited species, the greater prairie chicken. Near threatened, the greater prairie chicken has lost much of its habitat, with some populations experiencing extreme bottlenecks (37). To train our GWAC model, our neutral-enriched dataset consisted of all variants discovered from whole genome sequencing of four greater prairie chickens. Our functional-enriched variants were randomly generated. We used the same feature categories as in our human GWAC version described above. In total, the greater prairie chicken version of GWAC trained on nearly 22 million variants.

Feature weights found by the greater prairie chicken version of GWAC were similar to those found by the human version (Fig. 3A). The highest weighted feature was phyloP of 8 landfowl. This is consistent with what was found by our human version of GWAC, emphasizing that functional variation is associated with evolutionary conservation/acceleration in related species. Additional predictive features included genome context (CpG sites in 150bp window), further evolutionary conservation/acceleration scores, and variant attributes (e.g., reference allele). Notably, CpG sites were much more important in the greater prairie chicken version of GWAC. Additionally, SIFT scores were much less important in the greater prairie chicken version of GWAC. This difference may be caused by our use of SIFT scores that had relatively lower confidence for greater prairie chicken compared to human (see Methods). We note that Grantham distance was of increased importance in the greater prairie chicken version, possibly substituting for SIFT.

**Fig. 3.**
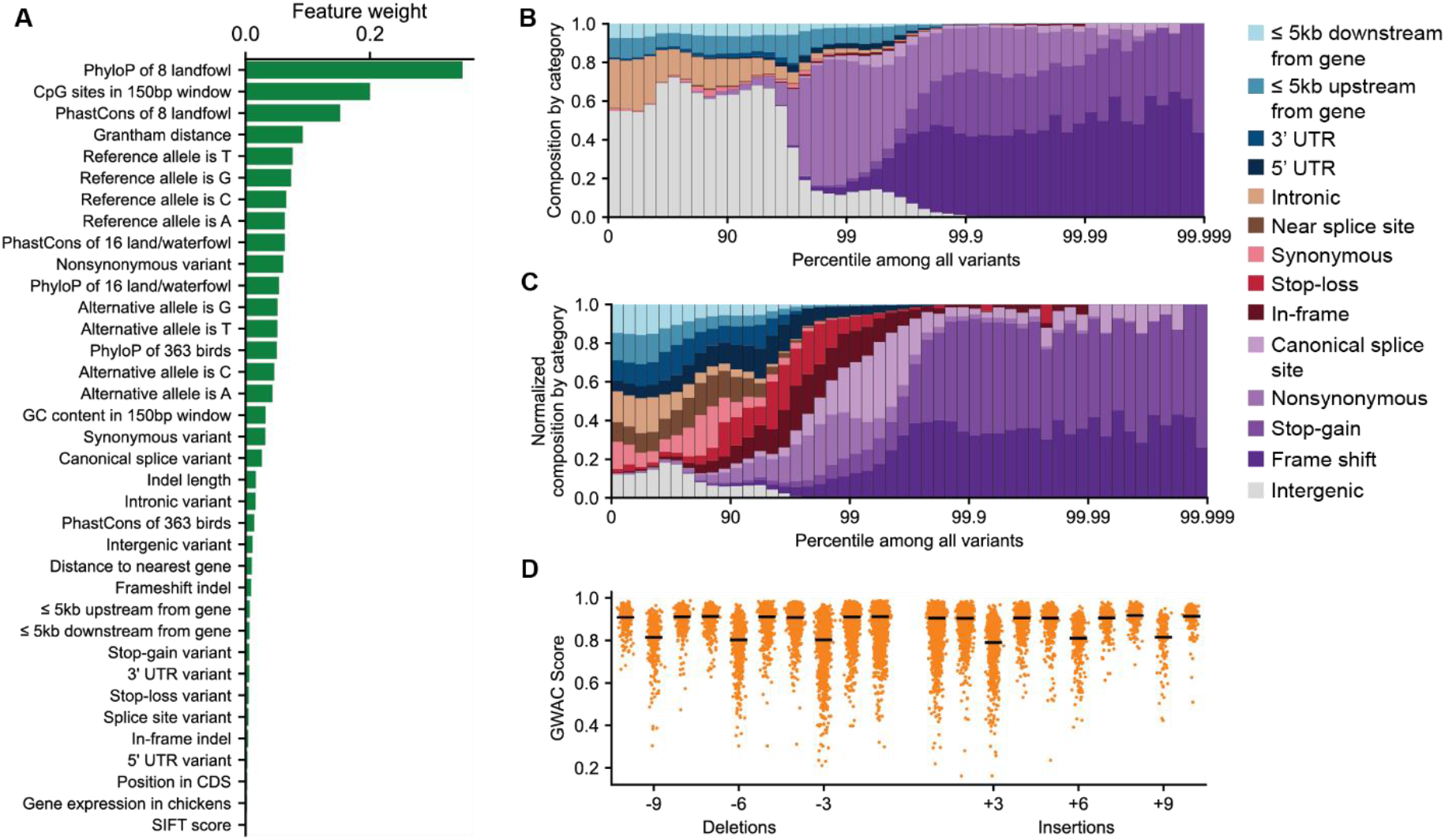
GWAC identifies general properties of functional variants in greater prairie chicken, consistent with our human version of GWAC. **A**) Absolute values of feature weights learned by GWAC. **B**) Composition of variant categories in each GWAC score percentile bin. 10 million variants were randomly generated to create this figure. Detailed descriptions of each consequence are provided in Table 1. **C**) Equivalent data as B, with variant categories normalized by the number of variants in each category. **D**) GWAC scores of frameshift and in-frame indels randomly generated in the greater prairie chicken genome. Black bars indicate mean values. Larger GWAC scores indicate greater chance of being functional.

The GWAC ranking of variants is comparable between the greater prairie chicken and human versions. Most variants in the 99^th^ percentile or above are frameshift, stop-gain, non-synonymous, or canonical splice site variants in the greater prairie chicken version (Fig. 3B,C), consistent with the human version. However, categories tend to span a larger percentile range in our greater prairie chicken version of GWAC. For example, variants above the 99.99 percentile were entirely frameshift variants in our human predictor. Yet, in our greater prairie chicken predictor, variants above the 99.99 percentile also consist of stop-gain, nonsense, and canonical splice site variants.

The greater prairie chicken version of GWAC identifies frameshift indels as more functionally important than in-frame indels, as does the human version. However, the difference in ranking between frameshift variants and in-frame variants is not as distinct as it was in the human version of GWAC. Additionally, the difference in ranking between insertions and deletions is much more subtle in our greater prairie chicken version of GWAC. It is difficult to know if these differences reflect true biological differences in indels between human and greater prairie chicken genomes, or chance differences in our models.

### Species-specific predictors retain variants that are lost during lift-over to model organism genomes

An appealing alternative to creating custom variant impact predictors for each individual species is to instead create variant impact predictors for well-characterized taxa, and then lift-over variants from nearby species to identify functional variants. This strategy has been used to prioritize variants in rhesus macaques with human CADD scores (38). To explore this possibility, we examined the correlation between variants ranked by different frameworks and species.

We used randomly generated variants to assess the degree of correlation between pairs of variant impact predictors. We found that GWAC and CADD scores were moderately correlated (Spearman’s *ρ* = 0.48) when evaluated on randomly generated variants in the human genome (Fig. 4A). The scores from these two predictors were most correlated on variants that each judged were highly likely to be functional (≥ 92^nd^ percentile). We next evaluated the correlation between GWAC and CADD-limited scores using the same variants, finding correlation was stronger (*ρ* = 0.64) and that variants judged to be either likely functional (≥ 92^nd^ percentile) or non-functional (≤ 4^th^ percentile) were most correlated (Fig. 4B).

**Fig. 4.**
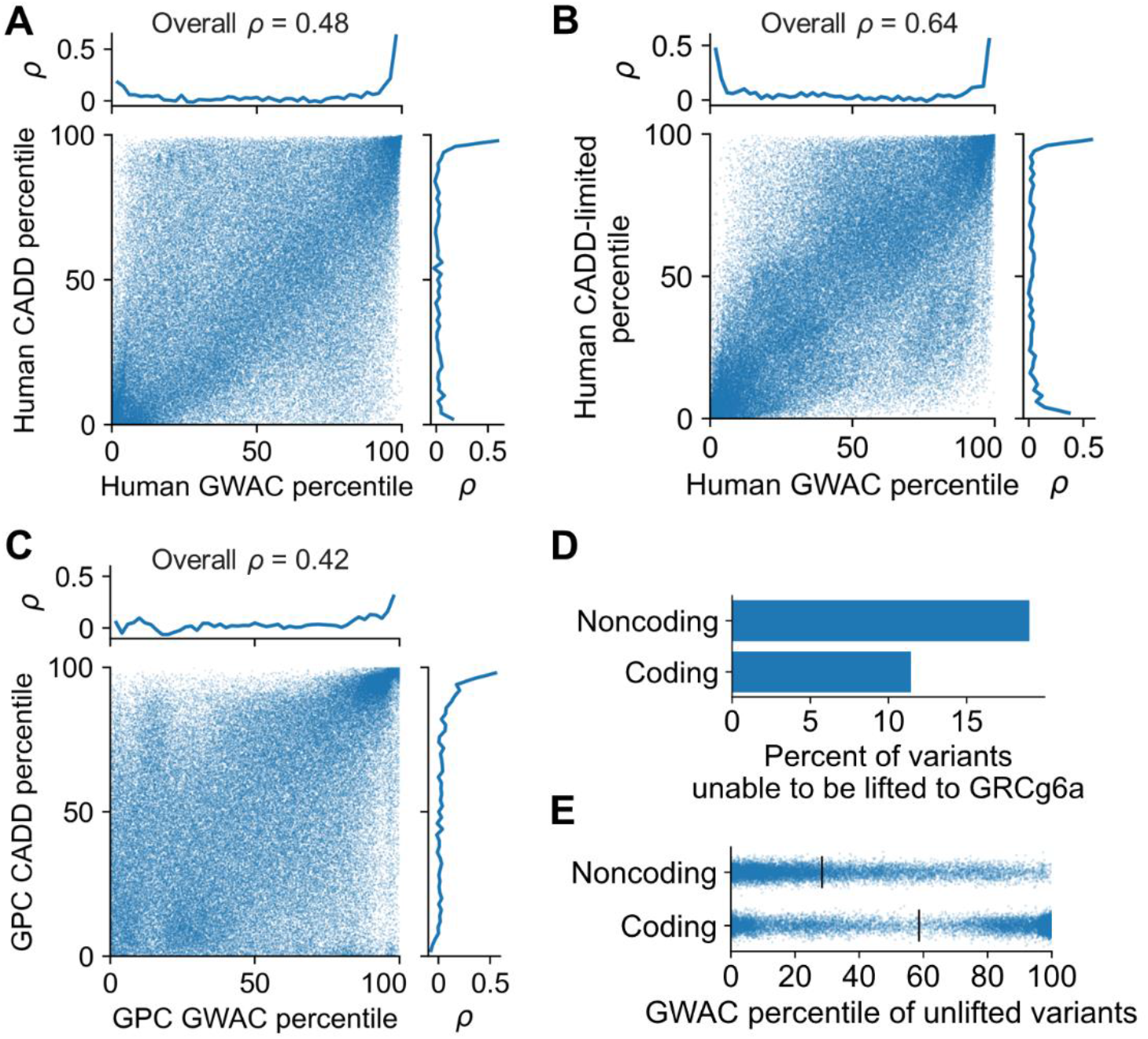
We find moderate correlation between methods, with particularly high correlation for variants in extreme percentiles. During the lift-over process, potentially functional variants are lost. **A**) Correlation of GWAC and CADD scores for 100,000 variants drawn randomly from the human genome. Marginal plots show Spearman’s rank correlation calculated in sliding windows. **B**) Equivalent analysis as in A) now considering GWAC and CADD-limited scores **C**) Equivalent analysis as in A) now considering greater prairie chicken GWAC and chCADD scores for 100,000 variants drawn randomly from the greater prairie chicken genome. **D**) Percent of random non-coding and coding variants that were unable to be lifted-over from the greater prairie chicken genome to GRCg6a. **E**) Scatter plot of GWAC percentile of variants that could not be lifted-over. Vertical black bars indicate mean values.

Next, we examined the correlation between GWAC and chCADD scores for variants randomly drawn from the greater prairie chicken genome. chCADD is a predictor developed for domestic chickens following the CADD framework. The domestic chicken is the closest species to the greater prairie chicken with an existing genome-wide predictor, although the two species diverged approximately 30-40 million years ago (39). After lifting-over variants from the greater prairie chicken genome to GRCg6a, we scored them with chCADD. We found that scores were moderately correlated (*ρ* = 0.42). Similar to our comparison of GWAC and CADD on human variants, we found that GWAC and chCADD scores were more correlated on variants judged by either to be likely functional (≥ 84^th^ percentile).

One barrier to the lift-over approach is that not all variants can be lifted-over to the target species. Among our random sample of variants in the greater prairie chicken genome, 11% of coding variants and 19% of non-coding variants were unable to be lifted-over to GRCg6a (Fig. 4D). Although it’s expected that variants in repetitive, poorly conserved, or low-complexity regions of the genome may not lift-over, we find that among unlifted coding variants, 30% have a GWAC score in the 95^th^ percentile or higher (Fig. 4E), suggesting these variants include a large fraction of potentially functional variants.

## Discussion

Here we’ve introduced GWAC, a method to differentiate functional from neutral variants in species with limited data. We’ve shown that from an annotated genome, we can predict functional variants with surprising accuracy. We evaluated a human version of GWAC, which was able to distinguish pathogenic from common variants nearly as well as CADD, which trains on dozens of additional features. When compared to SnpEff, a method commonly used to evaluate functional variants when only an annotated genome is available, GWAC made substantially better predictions for variants in several categories, such as missense or non-coding (Fig. 2C,D). When limited data are available, we found that the GWAC framework was always comparable or better than the CADD framework. Given the advantages of the GWAC framework (small sample sizes required, no ancestral genome needed), we anticipate that GWAC will be a more accessible framework for researchers to use when data is limited. We next developed a greater prairie chicken version of GWAC and found that it learns characteristics of functional variants similar to those learned by our human version of GWAC. Although it’s not possible to definitively evaluate our greater prairie chicken version of GWAC, this similarity to our human predictor suggests that it too can distinguish functional from neutral variants.

Additionally, we considered the trade-offs between generating new predictors for each species or lifting-over variants and using variant impact predictors in model organisms. The lower overall correlation between greater prairie chicken GWAC and chCADD (Fig. 4C) compared to human GWAC and CADD (Fig. 4A) could be interpreted as a small penalty for lifting-over variants between species. However, the largest drawback of lifting-over variants is that variants are lost during lift-over. We found that 11% of coding variants were lost when lifting-over variants from greater prairie chicken to chicken, and many of these variants were predicted to be functionally important. We expect that our ability to lift-over coordinates between greater prairie chicken and chicken is better than that of most non-avian species due to strong sequence and synteny conservation in birds (40). Thus, for many species of conservation interest, there will be even greater costs to lifting-over variants. Nonetheless, generating a new predictor does require considerable time and computational resources. We note that variant scores between the two approaches were most correlated for variants likely to be functional. Researchers interested in identifying functional variants with high specificity but not sensitivity may be best positioned to use the lift-over approach.

Our validation of GWAC in humans is limited by the constraints of ClinVar variants. Pathogenic variants in ClinVar are a biased sample of the total functional variants, reflecting those variants that have a sufficiently large effect to be associated with disease. These results may not accurately reflect a method’s ability to distinguish more moderately functional variants from neutral variants in a population.

We recognize that experimental validation of functional variants remains the gold standard for identifying variants with functional effects. Frameworks such as GWAC and CADD are not yet accurate enough for the predicted impact of a single variant to be high confidence. Despite these limitations, we see a vital role that in silico methods can play. For most species of conservation interest, there are limited experimental systems to assess variants. For extinct species, it may be impossible to experimentally validate variants in their original genetic context. In these cases, in silico predictors can be more rapidly developed to meet research questions. Even for species in which variants can be experimentally evaluated, in silico tools can play a valuable role in prioritizing variants. For example, a researcher that uses GWAC to identify the top 100 highest-impact variants in a genome may not discover 100 functional variants, but this subset will be significantly enriched for variants with functional effects.

We have taken care to use methods that are actively maintained and supported. This has required that we leave out some potentially valuable methods which are no longer maintained (GERP) or which have not been updated to run efficiently on large datasets (PROVEAN). Researchers willing to grapple with these challenges may find that incorporating these and similar methods improves performance.

Our work expands what was believed possible for functional variant prediction. Currently there are no genome-wide variant impact frameworks available for non-model organisms. We’ve demonstrated that large sample sizes and functional genomic annotations are not required for accurate predictions. Our findings are supported by previous work in human and mouse showing CADD prediction accuracy drops only modestly as functional genomics annotations are removed (21). Using the GWAC framework, researchers can now develop variant impact predictors for any species with an annotated genome and a few sequenced individuals. Our results suggest that these predictors will perform just slightly below the accuracy of one of the most widely used methods in humans.

Researchers often wish to distinguish between functional variants that are adaptive and functional variants that are deleterious. Although predictors such as GWAC can prioritize variants likely to be functional, the adaptiveness or deleteriousness of these variants can only be determined by measuring the fitness of organisms with these variants. However, we anticipate that GWAC can still be used to refine metrics of conservation interest. For example, GWAC scores should be lower in runs of homozygosity in species that have purged deleterious variants than in species that have not purged deleterious variants. Additionally, GWAC scores should be on average elevated in species with greater adaptive potential.

We envision that GWAC predictors can play a valuable role in conservation, particularly in applications where it is valuable to identify a subset of variants that are enriched for functional effect. For example, in species for which GWAS has been performed, variant impact predictors can help in fine-mapping, identifying the specific variants in GWAS peaks that are most likely causal. Similarly, GWAC can help prioritize variants for validation and potentially eliminate false positives from datasets during genotype-environment association analyses. Additionally, researchers may be interested in identifying specific variants that are responsible for differences between closely-related species. Even single variants may be responsible for important phenotypic differences between species, as illustrated by a single variant in NOVA1 distinguishing humans from Neanderthals in important brain regions (41). Identifying similar variants may be valuable to understand the genetic underpinnings of adaptation or to resurrect phenotypes lost to extinction (42).

Existing literature suggests evolutionary conservation/acceleration data are more useful for functional annotation of variants when they come from relatively small phylogenies of closely related species than large phylogenies that include more distant species (21, 26), and the feature weights learned by GWAC support these conclusions. This is likely explained by increasingly divergent selection on orthologous variants as species become more distant and their biological context changes. Most large-scale reference genome projects are focused on broadly sequencing species from as many orders as possible. While this strategy is valuable for characterizing overall diversity, gains for functional variant prediction will be greater from densely sequencing many species in the same family or genera, as this will permit the creation of multiple genome alignments from closely-related species. For example, predictions of functional missense variants in humans were recently improved by incorporating data from newly sequenced primate genomes (26).

Our goal was to determine if we could design a predictor to evaluate variants genome-wide within the constraint of reasonable computational resources and time. However, the resources and time required remain significant. The most time-consuming features to generate are the multiple genome alignments and their associated phyloP/phastCons scores. This burden will be reduced by continued developments to rapidly build multiple genome alignments (43) and compute conservation scores directly on these alignments (44). Nonetheless, for those with sufficient resources, it is likely that more computationally intensive methods will perform better, specifically for scoring missense variants. For example, EVE is an unsupervised deep learning method developed for humans that performs excellently on missense variants (45). EVE trains on a catalog of amino acid sequences from thousands of species, and thus a version of EVE could be generated for any sequenced species. However, EVE relies on training a separate neural network for each protein, and thus requires substantial computational resources that are currently beyond the scope of most academic labs.

Researchers have found that non-coding variation is responsible for substantial phenotypic variation both within and between species (46-48). Although GWAC performs better than random when identifying functional non-coding variants, they are the most challenging category to predict accurately. Even with rich functional genomics data, non-coding variant prediction remains a challenge for human predictors. The most accurate non-coding methods incorporate data from large-scale functional genomics experiments (e.g., histone modifications, open chromatin)(49-52), which will be challenging to carry out at scale in species of conservation interest. Promising methods to address these deficiencies may include machine learning methods that attempt to learn the language of genomic elements across species (53).

## Materials and Methods

### Data sources for the derived variant and common variant predictors in humans

For the common variant predictor, we sampled SNVs by retaining all variants identified in three randomly selected individuals in 1KGP (HG00157, HG01597, NA18853). We consider these sampled variants to be our ‘common’ variant set, but since we retained all variants, some small fraction will be rare. We generated simulated SNVs by following methods in Kircher et al (54). Briefly, we simulated variants using local mutation rate estimates in blocks of 100 kb. We estimated these local rates from our sampled variants, to ensure the simulated variants would match the distribution of the sampled variants. This simulation script is available at https://cadd.gs.washington.edu/simulator. Next, we annotated these variants with VEP (55) to determine variant consequence, SIFT, distance to nearest gene, amino acid change, and position in cDNA sequence. We used VEP’s custom annotation feature to determine variant scores for gerp++ of 100 vertebrates, 7-way phyloP, 20-way phyloP, 100-way phyloP, 7-way phastCons, 20-way phastCons, and 100-way phastCons. We downloaded these custom features from https://hgdownload.soe.ucsc.edu/goldenPath/hg38/. We annotated variants with CpG and GC content of the surrounding +/-75bp using custom scripts. Variants were annotated with the TPM (transcripts per million) of any gene in which they were contained, including both exons and introns. For variants overlapped by multiple genes, the largest TPM was used.

For the derived variant predictor, human-derived and simulated SNVs and indels were downloaded from the CADD v1.6 training data website (https://krishna.gs.washington.edu/download/CADD-development/v1.6/training_data/GRCh38/) in GRCh38 build. These variants were then annotated by VEP as described above. However, to preserve features that are unique to CADD (for example, conservation scores from multiple sequence alignments with human sequence removed) the following features were retained from the original CADD features provided by the CADD training data website: GC, CpG, cDNApos, SIFTval, priPhCons, mamPhCons, verPhCons, priPhyloP, mamPhyloP, verPhyloP, GerpS, Grantham.

In all predictor training data, only insertions or deletions of 50 or fewer basepairs were retained. Indels in which both the ref and alt allele were longer than 1bp were removed.

ClinVar variants were downloaded from the ClinVar FTP site on 29 Jan 2022 (an archived version is available here: https://ftp.ncbi.nlm.nih.gov/pub/clinvar/vcf_GRCh38/archive_2.0/2022/clinvar_20220129.vcf.gz). VEP was used to annotate the vcf with 1KGP variant frequencies. Cyvcf2 (56) was used to parse the vcf and retain only SNPs. Variants were retained if they had a review status of ‘criteria_provided,_single_submitter’, ‘criteria_provided,_multiple_submitters,_no_conflicts’, ‘reviewed_by_expert_panel’, or ‘practice_guideline’ AND a clinical significance of ‘Likely_benign’, ‘Benign’, ‘Pathogenic’, ‘Benign/Likely_benign’, ‘Likely_pathogenic’, or ‘Pathogenic/Likely_pathogenic’. Retained variants were then further filtered to remove variants with a 1KGP frequency greater than 1% in any of the 5 continental ancestries. ClinVar variants at the same position as variants in either the GWAC or CADD-like predictor training set were removed. The clinical significance for ClinVar variants was then binarized as either 1 (containing ‘pathogenic’) or 0 (containing ‘benign’). Only autosomal or X-chromosome variants were retained. To create our CADD-like predictor, these same ClinVar variants were annotated with features using the vcf upload tool at https://cadd.gs.washington.edu/score, with “GRCh38-v1.6” and “Include Annotations” selected.

### Training the derived variant and common variant predictors in humans

For each predictor we subsampled the simulated variants such that their distribution across chromosomes exactly matched that of the common/derived variants. GWAC was implemented as a logistic regression classifier in Python with scikit-learn v0.17 (57) with class LogisticRegression used to train each model. We performed a grid search to find the optimal hyperparameters by using a leave-one-chromosome-out cross validation strategy on the training data. Hyperparameters searched included the solver (‘lbfgs’, ‘liblinear’, ‘newton-cg’, ‘newton-cholesky’, ‘sag’, or ‘saga’), the penalty (‘l2’, ‘l1’, ‘elasticnet’, or None), and the c_value (100, 10, 1.0, 0.1, or 0.01). We found the following combination performed best: solver: ‘lbfgs’, penalty: ‘l2’, and c_value: 10.

### Data sources for the common variant predictor in greater prairie chicken

We sampled SNVs by retaining all variants identified in four randomly selected greater prairie chickens from Johnson et al. (60). We removed variants for which all sample genotypes were either homozygous reference or missing. We also removed variants for which all sample genotypes were either homozygous alternative or missing. Finally, we retained variants where at least one sample had a PHRED-scaled probability of homozygous reference >30 AND at least one sample had a PHRED-scaled probability of homozygous alternative >30 (i.e., at least one sample had a less than 1/1000 chance of being homozygous reference, and at least one sample had a less than 1/1000 chance of being homozygous alternative).

SIFT scores were calculated for the GPC genome by first creating a SIFT4G database following https://github.com/pauline-ng/SIFT4G_Create_Genomic_DB and then annotating the training variants and fixed differences by following https://github.com/pauline-ng/SIFT4G_Annotator (58).

### Comparison with chCADD

Fixed differences between the greater prairie chicken and heath hen were lifted-over from the greater prairie chicken genome to the galgal6 genome. Lift-over chain files were created using the following method: lastz was used to align each scaffold of the greater prairie chicken genome to the entire galgal6 genome, with the following parameters: hspthresh=22000, inner=2000, ydrop=3400, gappedthresh=10000, scores=bin/HoxD55, --chain, format=axt. axtChain was then used to chain together alignments, with the following options: - minscore=5000 -linearGap=loose. chainMergeSort was used to merge short chains, and chainSort was used to concatenate and sort all chains. chainNet was used to create nets from the sorted chains. Finally, netChainSubset was used to create a lift-over chain file from the nets. Picard LiftoverVcf was used to lift-over fixed differences to the galgal6 genome. Out of 364,957 variants, 39,772 (10%) failed to lift-over due to no target; 180,985 failed to lift-over due to mismatching reference alleles after lift-over (50%); and 144,200 (40%) successfully lifted-over. The large number of variants that failed to lift-over due to mismatching reference alleles after lift-over is likely due to the fact that we are selecting for those positions that are recently derived. Since chCADD only scores variants on chromosomes 1-33 and Z, we removed variants on other chromosomes.

### Generation of multiple genome alignment and associated features

The greater prairie chicken (GRPC) genome (GCA_001870855.1) was improved using Satsuma v. 3.1.0 (59) as described in Johnson et al. (60). The 363-way avian Cactus alignment was downloaded from https://cgl.gi.ucsc.edu/data/cactus/363-avian-2020.hal. To replace the unmodified GRPC genome in the hal file with the improved GRPC genome, the following steps were taken. halRemoveGenome (https://github.com/ComparativeGenomicsToolkit/hal) was used to remove the unmodified GRPC genome from the hal file as well as the turkey genome, as GRPC and turkey are sibling leaf nodes in the hal alignment. The ancestral genome of turkey and GRPC was retrieved from the hal file using hal2fasta. Cactus was used to align the ancestral genome, the turkey genome, and the improved GRPC genome. The function halAppendSubtree was used to merge this subtree to the larger hal alignment, resulting in a 363-way alignment with the improved GRPC.

Hal2mafMP (https://github.com/ComparativeGenomicsToolkit/hal/blob/master/maf/hal2mafMP.py) was used to convert the hal alignment to a maf file for each scaffold. Due to repetitive regions in genomes, some aligned regions in the maf files contained multiple sequences from the same species. Maf_stream (https://github.com/joelarmstrong/maf_stream) was used to identify the consensus sequence for each species, and output a maf file with no more than one sequence from each species per region.

PhyloP from PHAST (11) was used with the options “--method LRT --mode CONACC --wig-scores” to generate per-base phyloP scores. phastCons from PHAST was used with the options “--expected-length=45 --target-coverage=0.3 --rho=0.3” to generate per-base phastCons scores. Three levels of phylogeny were considered. The broadest level considered all 363 birds in the alignment. The middle level considered all landfowl and waterfowl (16 genomes). The narrowest considered only landfowl (8 genomes). For the 8 and 16-bird alignments, Maffilter (https://bmcgenomics.biomedcentral.com/articles/10.1186/1471-2164-15-53) was used to subset the maf files to include only the species of interest.

## Acknowledgements

The author would like to thank Jeff A. Johnson, Ben. J. Novak, Shamil R. Sunyaev, Nilah M. Ioannidis, and Beth Shapiro for their contributions to this work. The author would also like to thank Daren Card for valuable computational advice as well as Adam Siepel and Rebecca Hassett for addressing an underflow issue in phast. AS was supported by an NSF PRFB Program under Grant No. 2109912.

