## Supplementary figures for "GWAC: A machine learning method to identify functional variants in data-constrained species"

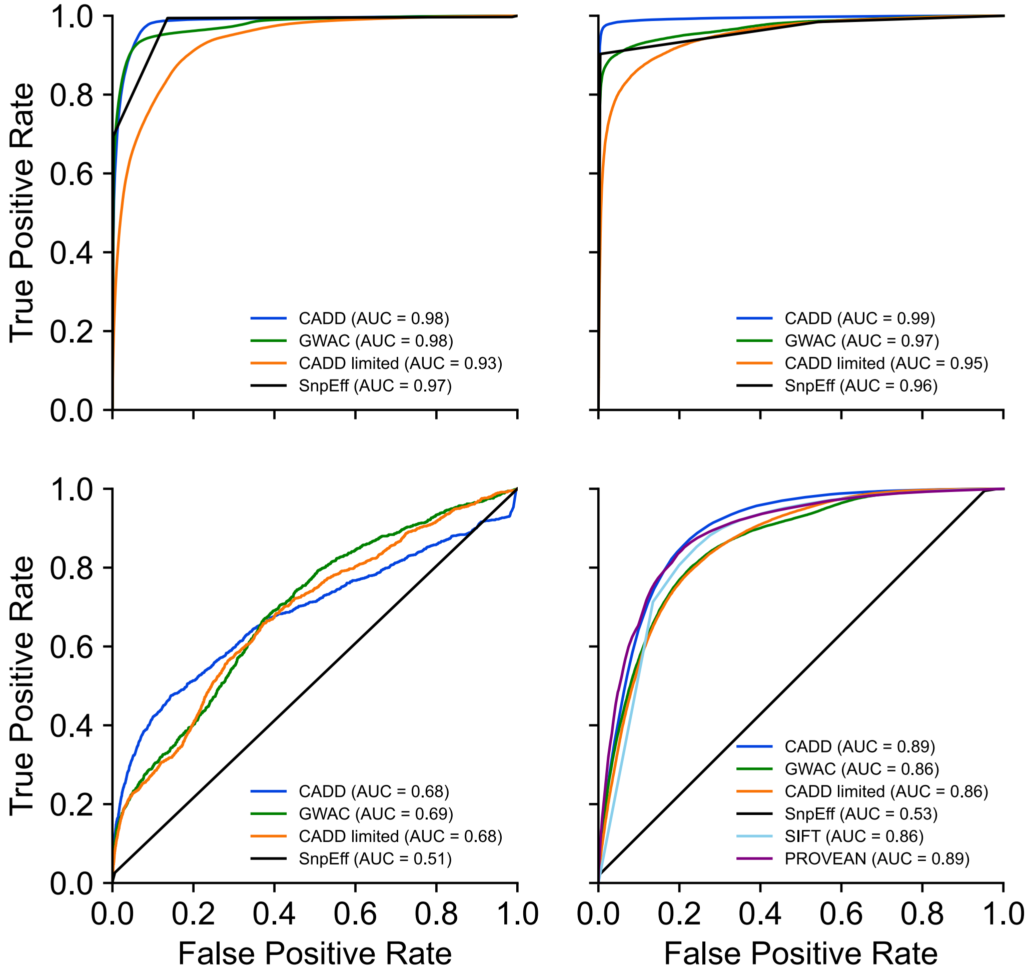


**Fig. S1** This figure is an equivalent analysis to Fig. 2 but is instead showing receiver-operating characteristic (ROC) comparison of GWAC with other genome-wide predictors on ability to distinguish pathogenic from *rare benign* variants in **A**) coding regions **B**) splicing regions, and **C**) non-coding regions. **D**) ROC comparison of GWAC with missense predictors. Rare benign variants were collected from ClinVar.
